# Sex- and caste-specific developmental responses to juvenile hormone in an ant with maternal caste determination

**DOI:** 10.1101/2023.09.29.559719

**Authors:** J. Brülhart, A. Süß, J. Oettler, J. Heinze, E. Schultner

## Abstract

Queen-worker caste polyphenism in social insects is a prime example for developmental plasticity. Most of what we know about caste development comes from studies of the honeybee, in which female caste is determined during larval development and workers retain functional ovaries. The ant genus *Cardiocondyla* is one of only few genera in which complete worker sterility has evolved, so that adult workers completely lack reproductive organs. In *C. obscurior*, queen- and worker-destined individuals are distinct in their development by late-embryogenesis, and castes can be distinguished in a non-invasive manner from this stage onwards. This provides the opportunity to investigate the degree of flexibility in caste development in a species with early caste determination. Using topical juvenile hormone treatment, a method known to influence caste determination and differentiation in some species, we investigated whether hormone manipulation affects the development and growth of queen and worker-destined late-stage embryos and larvae, as well as of early-stage embryos which cannot yet be distinguished by caste. We found no effect of hormone treatment on female caste ratios or body sizes in any of the treated stages, even though individuals reacted to heightened hormone availability with increases in the expression of *krüppel-homolog 1*, a conserved JH first-response gene. In contrast, hormone treatment resulted in the emergence of significantly larger males. These results show that in *C. obscurior*, early, presumably maternal caste determination leads to irreversible and highly-canalized caste-specific development and growth.

## Introduction

Almost 50 years ago, juvenile hormone (JH) was proposed to be a master regulator of social insect polyphenism (Nijhout & Wheeler, 1982; Wheeler, 1986, 1991). For ants, this was based on studies of soldier caste determination in *Pheidole bicarinata* (Wheeler & Nijhout, 1981, 1983), and a study in *Pheidole pallidula* which demonstrated that increasing JH levels in queens and eggs could lead to the production of new queens (Passera & Suzzoni, 1979). It had also been shown that JH treatment of late instar *Myrmica rubra* larvae could induce queen development, provided these larvae had overwintered and reached a critical size (Brian, 1974). Finally, JH treatment of first and second instar *Solenopsis invicta* larvae, but not queens or eggs, had been reported to result in increased queen production (Vinson & Robeau, 1974; Robeau & Vinson, 1976; Banks *et al*., 1978). Analogous to results on JH, differences in ecdysteroid titers had been detected in queen- and worker-destined eggs and larvae (Suzzoni *et al*., 1980, 1983).

Since Wheeler’s seminal work, to the best of our knowledge, five additional studies have investigated the relationship between hormone signalling and queen-worker development in ants (Schrempf & Heinze, 2006; Cahan *et al*., 2011; Penick *et al*., 2012; Libbrecht *et al*., 2013; Kuhn *et al*., 2018), and only one has followed up on previous reports (de Menten *et al*., 2005), so that empirical evidence for the generality of JH-regulation of caste and caste-specific body size is still surprisingly limited. One reason for this is the difficulty of studying development in ants: many do not produce sexuals in the lab and, unlike honey bees, ants do not rear castes in distinct brood cells, making it challenging to follow individual development and near impossible to identify castes before size and shape differences appear in late-stage larvae (Schultner & Pulliainen, 2020).

The genus *Cardiocondyla* is one of only few ant genera (ca. 3% of ∼300 genera) in which complete worker sterility has evolved, meaning adult workers lack reproductive organs. We recently showed that in *C. obscurior*, queen- and worker-destined embryos and larvae can be distinguished in a non-invasive manner using caste-specific crystalline deposits (Schultner *et al*., 2023). Queens appear to have full control over the production of daughter queens, which increases as queens age (Jaimes-Nino *et al*., 2022), indicating that female caste is determined via maternal effects in this species (Schultner *et al*., 2023). In addition to the two female castes, *C. obscurior* also produces two discrete male morphs – large, winged males and small, wingless males (Kinomura & Yamauchi, 1987; Heinze & Hölldobler, 1993; Oettler *et al*., 2010). When male morph is determined is unknown, but it is presumed to occur later than in females (Schrempf & Heinze, 2006). This unique system allowed us to test 1) how a presumably maternal mode of female caste determination, associated with early morphological differentiation, affects the degree of flexibility in caste-specific development and 2) whether male and female polyphenism are regulated in a similar manner. We did this by subjecting embryos and larvae to juvenile hormone treatment, which has previously been linked to increased queen and winged male production in *C. obscurior* (Schrempf & Heinze, 2006). We documented effects on adult caste and morph ratios and on larva and adult body sizes, as well as on the expression of the conserved JH-response gene *krüppel-homolog 1*.

## Results and discussion

### Hormone treatment does not affect caste or morph ratios in C. obscurior

A previous study reported that treatment of *C. obscurior* larvae of unknown sex and caste with the synthetic JH-analogue methoprene always resulted in the production of winged morphs, i.e. queens and winged males (Schrempf & Heinze, 2006). We first attempted to replicate this result by treating larvae which were collected from stock colonies without regard to caste-specific crystalline deposits. Compared to a solvent control, methoprene treatment (2 µl of a 1 mg/ml solution in 70% ethanol) did not have a significant influence on winged morph production in any of the three larval stages (Fisher’s exact test, ethanol solvent control vs. methoprene: first instar larvae: p=0.241, second instar larvae: p=0.573, third instar larvae: p=0.86; Table S1).

Similarly, experimental hormone treatment had no effect on the caste of emerging females when queen and worker-destined late-stage embryos and larvae were separated according to crystalline deposit patterns and treated separately (Table 1). Across all developmental stages and treatments, presumed queen-destined individuals developed into adult queens in 87% (346/397) of cases, and presumed worker-destined individuals developed into workers in 97% (321/331) of cases. In the three larval stages, the accuracy of caste fate prediction was >90% in individuals subjected to a handling control, and this remained similar following treatment with a solvent control (1 µl of 70% ethanol), as well as after treatment with JH III or methoprene (1 µl of a 1 mg/ml solution in 70% ethanol, Table 1). We confirmed the irreversibility of caste determination by treating queen- and worker destined second instar larvae with higher doses of hormone (2 µl of 1 mg/ml JH and methoprene); in all cases, treated individuals developed into the predicted caste (Table S1). Compared to larvae, queen caste was more difficult to predict in eggs, with only around half of untreated, presumed queen-destined eggs developing into queens; this proportion did not change after treatment (Table 1). In contrast, all untreated, presumed worker-destined eggs developed into workers, indicating that worker-destined eggs are more easily identified than queen-destined eggs and/or more likely to end up in experiments because they outnumber queen-destined eggs in stock colonies. The accuracy of caste prediction in early developmental stages furthermore appears to be influenced by sampling precision, as a previous study showed higher prediction accuracy rates for queen-destined eggs, but lower rates for queen-destined first instar larvae (Schultner et al. 2023). Caste prediction accuracy was lowest for methoprene-treated worker-destined eggs, but still relatively high at 80% (12/15), suggesting minor, if any, effects of methoprene on caste once it can be identified by caste-specific crystalline deposits. Of the presumed queen- and worker-destined individuals which emerged as adults, 4.1% (31/759) developed into males, the majority in replicates with presumed worker-destined individuals. Male morph fate was not affected by treatment, as only wingless males emerged (Table 1).

**Table 1:**
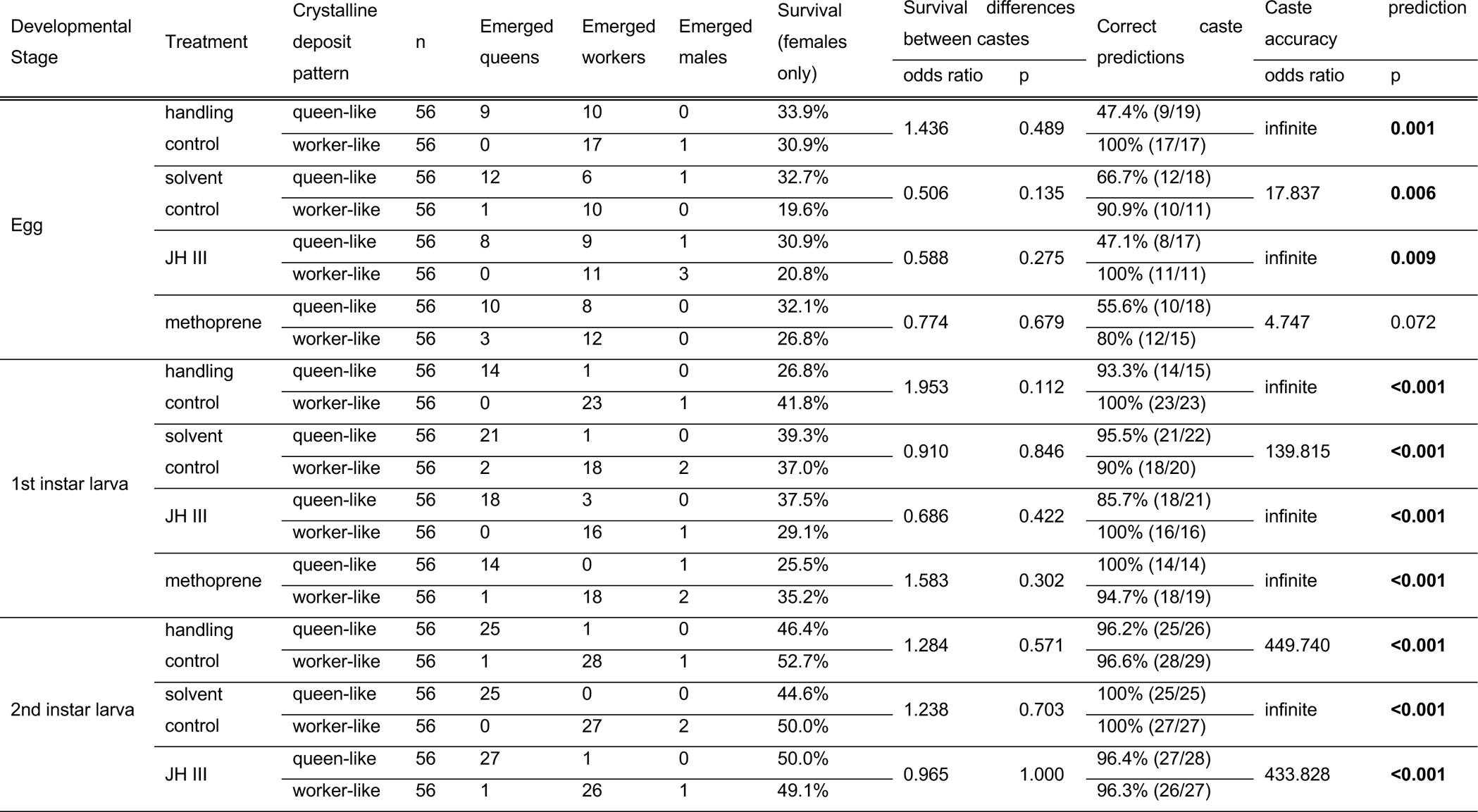
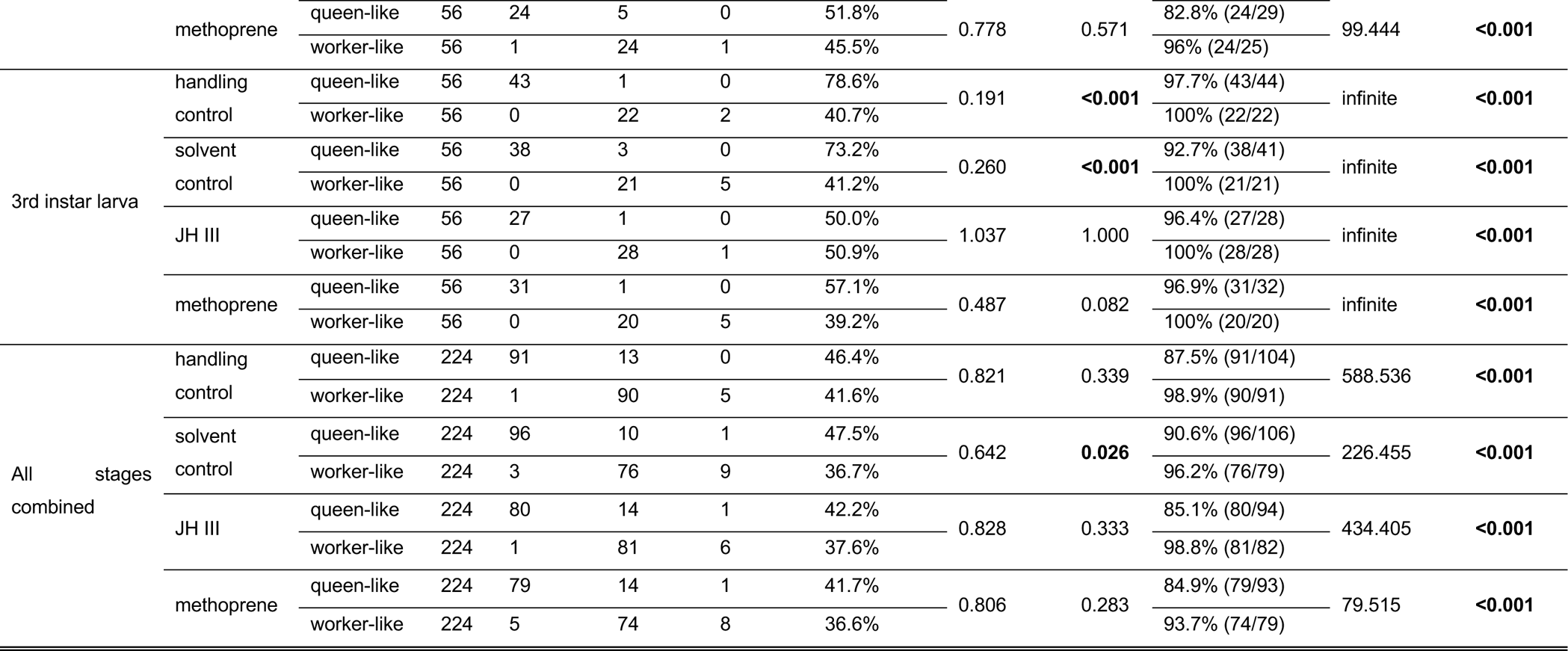
Survival and caste ratios of queen- and worker-destined embryos and larvae after hormone treatment.

Caste ratios were also not affected by treatment of female early-stage embryos (1-24 hours old), with 2.9% (1/35) of solvent control-treated eggs developing into queens, compared to 7% (4/57) of methoprene-treated eggs (Fisher’s exact test, odds ratio=0.393, p=0.646). Methoprene-treated eggs showed slightly higher survival (methoprene: 20% (57/285), solvent control: 12.3% (35/285), Fisher’s exact test, odds ratio=1.784, p=0.016). When early-stage male embryos produced by unmated queens were treated, only wingless males emerged in both treatments (ethanol: 100% (10/10), methoprene: 100% (11/11), Fisher’s exact test: odds ratio=infinite, p=1), and treatment did not affect male survival, which was extremely low (ethanol: 3.1% (10/325), methoprene: 3.4% (11/325), Fisher’s exact test: odds ratio=1.103, p=1).

Juvenile hormone analogues such as methoprene can be used to kill insects because high doses of these chemicals perturb development (Troisi & Riddiford, 1974; Tay & Lee, 2014; Nur Aliah *et al*., 2021). Similarly, in *C. obscurior*, survival of treated brood dropped considerably when high doses (5 mg/ml) were administered (Schrempf & Heinze, 2006). The moderate dose used here did not have adverse effects on survival in queen- and worker-destined individuals, which was similar between treatments in each of the developmental stages (Table 1). Across all developmental stages and treatments, a higher proportion of queen-destined compared to worker-destined individuals survived until pupation (queen-destined: 44.5% (346/893), worker-destined: 38.1% (321/868); Fisher’s exact test, odds ratio=0.770, p=0.008); this was mostly driven by differential survival of castes in the third larval instar (Table 1). Adult workers do not discriminate between developing queens and workers (Schultner *et al*., 2023), so differential treatment is unlikely to explain this difference. Instead, increased survival of queen-destined larvae may stem from size differences between the two castes (Oettler *et al*., 2019; Schultner *et al*., 2023).

### Male but not female body size is affected by hormone treatment

Ant body size is strongly associated with caste polyphenism, with queens typically larger than workers (see (Trible & Kronauer, 2017, 2021a, 2021b; Abouheif, 2021) for a discussion about the role of body size in ant caste development). Diverse factors have been shown to be associated with body size variation in queens and workers, including genotype, maternal effects, nutrition, as well as social and abiotic environment, e.g. (Heinze *et al*., 2003; Hughes *et al*., 2003; Bargum *et al*., 2004; Fjerdingstad, 2005; Schwander *et al*., 2005; Meunier & Chapuisat, 2009; Linksvayer *et al*., 2011). How caste-specific queen and worker body sizes are attained is not well understood, but studies focusing on size variation within the worker caste have brought valuable insight. Methylation of *epidermal growth factor receptor*, which links to JH via the insulin signalling pathway, appears to play a key role in determining worker size, both in species with distinct worker castes (Alvarado *et al*., 2015) and in species with monomorphic workers (Renard *et al*., 2022). JH itself has been implicated in increased body size of *Pheidole* soldiers (Wheeler & Nijhout, 1981, 1983; Rajakumar *et al*., 2012) and *Pogonomyrmex* and *Camponotus* workers (Cahan *et al*., 2011; LeBoeuf *et al*., 2016).

In *C. obscurior*, hormone manipulations did not result in larger queens or workers. After treatment in the second larval stage, queen-destined third instar larvae were always larger than worker-destined third instar larvae (Figure 1, Figure S1, Table 2, Table S2), confirming previous results (Schultner *et al*., 2023). Treatment had no effect on the size of worker larvae. Methoprene-treated queen-destined larvae were smaller than those subjected to a handling control, but there was no difference between any of the other treatments (Figure 1, Figure S1, Table 2, Table S2).

**Figure 1:**
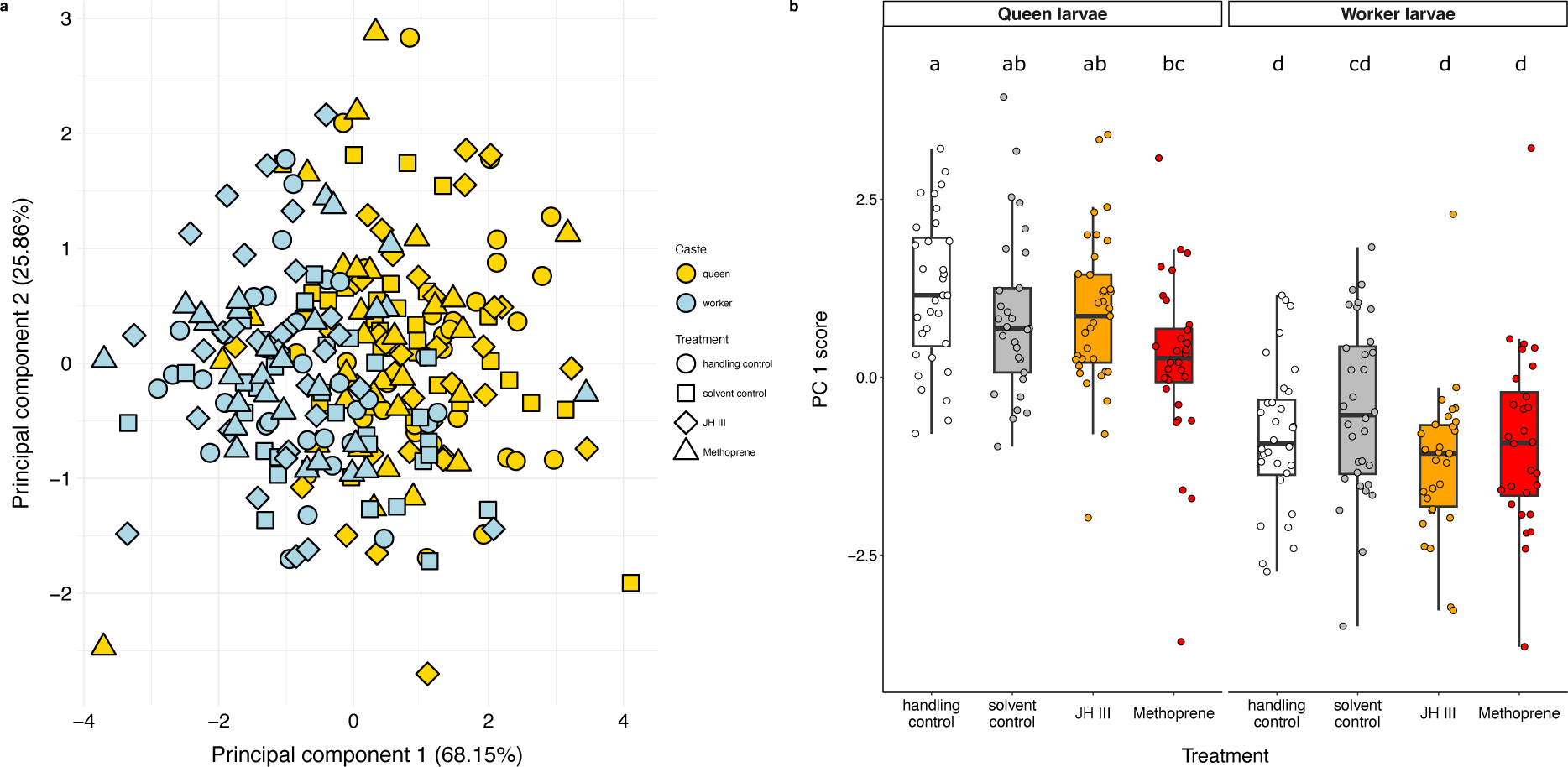
Body size of queen- and worker-destined larvae after hormone treatment. a) Principal component analysis of larval head width, body width and body length separates larvae by overall size on PC1. See table S3 for loadings. b) Queen larvae are larger than worker larvae across treatments. See table S2 for Tukey-corrected pairwise p-values and figure S1 for boxplots of individual traits.

**Table 2:** Effects of hormone treatment on body size and development time in the ant *Cardiocondyla obscurior*.

| Trait | Developmental stage treated | n | Linear regression |  | df | Sum of squares | F/X<sup>2</sup> | p |
| --- | --- | --- | --- | --- | --- | --- | --- | --- |
| Larva size | L2 larva | 250 | PC 1 score ~ | caste | 1 | 168.188 | 128.917 | **<0.001** |
|  |  |  |  | treatment | 3 | 13.201 | 3.373 | **0.019** |
|  |  |  |  | caste * treatment | 6 | 12.007 | 3.068 | **0.029** |
| Adult queen, worker and male size | late-stage embryo, L1, L2 & L3 larva | 759 | PC 1 score ~ | caste | 2 | 1747.37 | 709.367 | **<0.001** |
|  |  |  |  | treatment | 3 | 27.84 | 7.534 | **<0.001** |
|  |  |  |  | developmental stage | 3 | 7.35 | 1.99 | 0.114 |
|  |  |  |  | caste * treatment | 6 | 38.4 | 5.196 | **<0.001** |
| Adult queen, worker and male size | early-stage male and female embryo | 74 | PC 1 score ~ | caste | 2 | 108.528 | 41.582 | **<0.001** |
|  |  |  |  | treatment | 1 | 3.392 | 2.599 | 0.112 |
|  |  |  |  | caste * treatment | 2 | 3.814 | 1.461 | 0.239 |
| Adult queen, worker and male size | early-stage male and female embryo | 74 | PC 2 score ~ | caste | 2 | 18.200 | 14.495 | **<0.001** |
|  |  |  |  | treatment | 1 | 2.345 | 3.736 | 0.057 |
|  |  |  |  | caste * treatment | 2 | 5.806 | 4.624 | **0.013** |
| Development time | late-stage embryo | 62 | days to pupation ~ | caste | 1 | 69.5 | 4.517 | **0.038** |
|  |  |  |  | treatment | 3 | 23.88 | 0.517 | 0.672 |
|  | L1 larva | 46 | days to pupation ~ | caste | 1 | 35.61 | 1.437 | 0.238 |
|  |  |  |  | treatment | 3 | 21.43 | 0.288 | 0.834 |
|  | L2 larva | 55 | days to pupation ~ | caste | 1 | 144.24 | 16.331 | **<0.001** |
|  |  |  |  | treatment | 3 | 62.98 | 2.377 | 0.081 |
|  | L3* | 58 | days to pupation ~ | caste | 1 | - | 0.704 | 0.401 |
|  |  |  | days to pupation ~ | treatment | 3 | - | 4.122 | 0.248 |

\*For each trait, data were analysed using a single linear regression except for development time of L3 larvae, which was compared between castes and treatments using separate Kruskal-Wallis tests as the data did not follow a normal distribution.

Adult queens emerging from treatments of late-stage embryos and larvae were also larger than adult workers (Figure 2, Figure S2, Table 2, Table S4). This was independent of the timing of hormone treatment, i.e. the developmental stage at which individuals were treated, so that data were pooled across developmental stages for subsequent analyses (Table 2). As in larvae, treatment did not result in larger queens or workers (Figure 2, Figure S2, Table 2, Table S4). Similarly, treatment had no clear effect on development time until pupation in any of the four treated stages (Table 2). Caste affected development time of late-stage embryos and second instar larvae, with queens exhibiting longer development times than workers; there was no effect of caste on development time of first and third instar larvae (Table 2).

**Figure 2:**
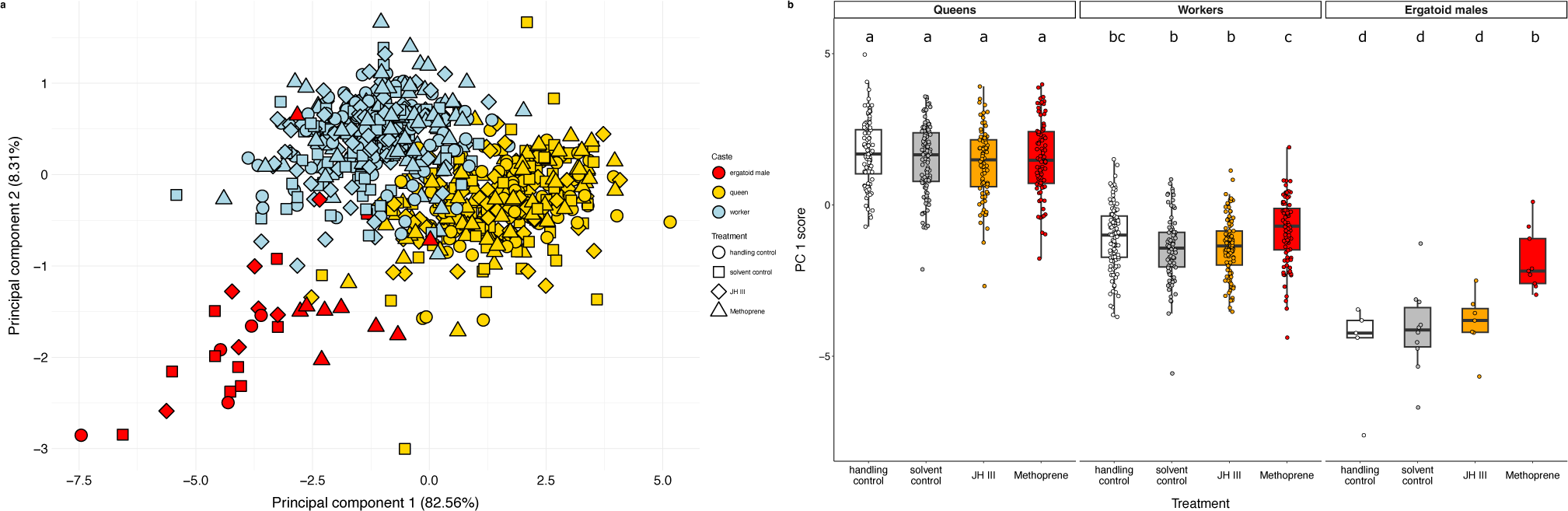
Body size of adult queens, workers and males emerging from hormone treatment of late-stage embryos and larvae. a) Principal component analysis separates individuals by size on PC1. See table S5 for loadings. b) PC 1 scores plotted by caste and treatment show that queens are the largest caste, and workers are larger than males. Female size is not affected by hormone treatment, whereas larger males emerge from methoprene treatments. See table S4 for Tukey-corrected pairwise p-values and figure S2 for boxplots of individual traits.

In contrast to females, males treated with methoprene emerged as significantly larger adults than males emerging from the other treatments (Figure 2, Figure S2, Table 2, Table S4). Wingless males are typically smaller than females (Oettler *et al*., 2019), but males emerging from methoprene treatments exhibited body sizes similar to those of workers (Figure 2, Figure S2, Table 2, Table S4).

Adult queens emerging from treatment of female early-stage embryos of unknown caste were larger than workers and males, and again, methoprene did not affect the size of workers or queens (Figure S4, Figure S5, Table 2, Table S6). Male morphology did respond to treatment of early-stage embryos, but in an unexpected manner: males emerging from methoprene treatments tended to exhibit shorter thorax lengths and petiole widths compared to solvent-control treated males (Figure S5, Table 2, Table S6).

### Conserved expression of a JH first response gene in queens, workers and males

Molecular pathways associated with juvenile hormone have been extensively studied in a few model organisms such as the tobacco hornworm *Manduca sexta* and the flour beetle *Tribolium castaneum* (Riddiford, 2012). These have a revealed a suite of conserved genes acting downstream of JH, including *methoprene-tolerant* and *krüppel-homolog* 1 (*Kr-h1*) (Konopova & Jindra, 2007; Minakuchi *et al*., 2008, 2009; Jindra *et al*., 2013). *Kr-h1* expression in social Hymenoptera has mainly been studied in adults, and varies e.g. with task in bee workers (Whitfield *et al*., 2003; Grozinger & Robinson, 2007; Shpigler *et al*., 2010; Johnson & Jasper, 2016), and with caste (Glastad *et al*., 2021; Zhang *et al*., 2021) and reproductive status in ants (Araki *et al*., 2020; Gospocic *et al*., 2021). During worker development, pharmacologically-induced hypomethylation resulting in increased expression of the growth-promoting *egfr* gene was associated with decreased *kr-h1* expression, and an increase in body size (Renard *et al*., 2022). Similarly, knockout of an estrogen-related receptor resulted in decreased *kr-h1* expression (Zhang *et al*., 2021). While these studies have helped confirm the molecular links between insulin and hormone signalling, growth and reproduction, it remains unknown whether the function of *kr-h1* as a JH first-response gene is conserved across sexes and castes in developing ants. We used quantitative PCR to measure *kr-h1* expression following hormone treatment in *C. obscurior*. In female larvae, *kr-h1* expression increased significantly 6 and 24 hours after hormone treatment compared to a solvent control (Figure 3, Linear regression, factor: treatment, 1h: F_2,39_=0.661, p=0.522; 2h: F_2,29_=2.044, p=0.144; 6h: F_2,38_=3.997, p=0.027; 24h: F_2,40_=10.016, p<0.001). In males, hormone treatment led to increased *kr-h1* expression after 1, 2 and 24 hours (Figure 3, Linear regression, factor: treatment, 1h: F_1,24_=6.767, p=0.016; 2h: F_1,23_=15.993, p<0.001; 6h: F_2,12_=1.27, p=0.282; 24h: F_1,23_=4.665, p=0.041).

**Figure 3:**
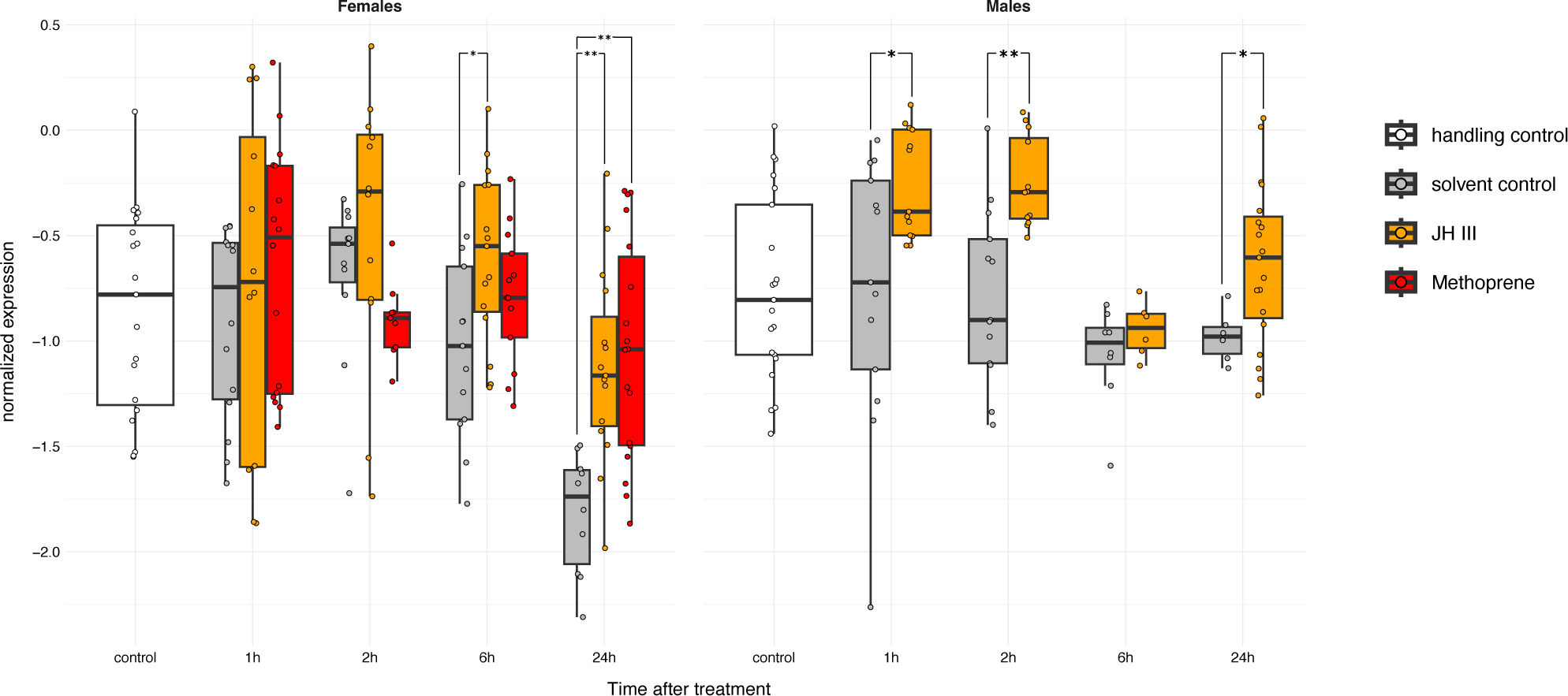
Timeline of *krüppel-homolog 1* expression in female and male larvae after hormone treatment. Expression in larvae from the three larval instars was compared between treatments, separately for each sex and timepoint, using linear regressions followed by Tukey-correction of p-values (*p<0.05, **p<0.01).

*Krüppel-homolog 1* expression was not affected by the caste of late-stage embryos or larvae (Figure S6, Linear regression, factor: caste, eggs: F_1,77_=0.045, p=0.833; L1 larvae: F_1,54_=3.785, p=0.057; L3 larvae: F_1,42_=0.035, p=0.853; L2 larvae: as model residuals did not fit assumptions of normality linear regressions comparing caste-specific expression were run separately by treatment, handling control: F_1,8_=0.033, p=0.86, solvent control: F_1,12_=0.188, p=0.672, JH III: F_1,13_=4.139, p=0.063, methoprene: F_1,10_=0.032, p=0.862). There was furthermore no effect of treatment in late-stage embryos, first instar and second instar larvae (Figure S6, Linear regression, factor: treatment, eggs: F_3,77_=0.522, p=0.668; L1 larvae: F_3,54_=1.934, p=0.135; L2 larvae: as model residuals did not fit assumptions of normality, linear regressions comparing expression by treatment were run separately in each caste, queens: F_3,23_=1.392, p=0.86, workers: F_1,20_=3.016, p=0.054). Third instar larvae of both castes exhibited significantly higher *kr-h1* expression after JH III treatment compared to handling control and solvent control-treated larvae (Figure S6, Linear regression, factor: treatment, L3 larvae: F_3,42_=5.574, p=0.002; see table S8 for Tukey-corrected pairwise p-values).

## Conclusions

Juvenile hormone studies in ants have employed a number of different methods, from topical application with fine capillaries or brushes to supplementation of liquid and solid food, to physical contact with hormone-soaked objects. In addition, a wide range of doses have been used and sometimes only cryptically reported, making it difficult to compare between studies. Furthermore, mortality was not always reported accurately, or controls were missing. This is further complicated by the range of factors associated with caste determination in ants, including genotype, maternal effects, nutrition, social environment, temperature and combinations thereof, e.g. (Brian, 1975; Passera, 1980; Helms-Cahan *et al*., 2002; Anderson *et al*., 2008; Smith *et al*., 2008; Penick & Liebig, 2012), and the consequences for the timing of determination. Accordingly, the interpretation of experimental studies on the role of JH in ant queen-worker caste development is not straightforward.

In *Cataglyphis mauritanica* and *Pogonomyrmex barbatus x rugosus*, two species with a genetic component to caste determination (Helms-Cahan *et al*., 2002; Julian *et al*., 2002), methoprene treatment of eggs (*C. mauritanica,* Kuhn et al. 2018) but not queens (*P. barbatus x rugosus,* Cahan et al. 2011) increased production of queens with non-hybrid (i.e., worker) genotypes. Colonies with treated *P. barbatus x rugosus* queens produced larger workers, but this presumed hormone effect cannot be disentangled from treatment-induced changes to colony size (Cahan *et al*., 2011). In *Pheidole pallidula*, only eggs produced by overwintered queens shortly after hibernation can give rise to new queens (Passera, 1980); treating queens and eggs with JH I outside this time period resulted in queen production, whereas treatment of larvae had no effect (Passera & Suzzoni, 1979). In another *Pheidole* species, methoprene treatment of queens, eggs and larvae did not result in queen production (Ono, 1982). In *Pogonomyrmex rugosus*, as in *P. pallidula,* only hibernated queens are capable of producing queen-potent eggs (Schwander *et al*., 2008); here, feeding hibernated colonies with methoprene strongly increased the proportion of produced queens (Libbrecht *et al*., 2013), though colony productivity and brood mortality were not monitored. In *Solenopsis invicta*, treatment with JH analogues of larvae but not eggs or queens resulted in queen production (Vinson & Robeau, 1974; Robeau & Vinson, 1976; Banks *et al*., 1978). Finally, in three species with caste determination during larval development, topical treatment with JH analogues resulted in development of more and larger queens in *M. rubra* (Brian, 1974) and increased queen production in *Harpegnathos saltator* (Penick *et al*., 2012), but feeding with JH III had no effect on queen production in *Camponotus floridanus*, though worker size increased (LeBoeuf *et al*., 2016).

How does *C. obscurior* fit into this admittedly confusing picture? Like in *P. pallidula* and *P. rugosus*, queen-potency of eggs is controlled by mother queens. However, being a tropical species, *C. obscurior* does not undergo hibernation, and queens are capable of producing queen-destined eggs at any time, though their proportions increase when queens become older (Jaimes-Nino *et al*., 2022). The doses we used here (between ca. 400 - 40000 µg/g body weight depending on the experiment) are within the general range reported in other studies (between ca. 60 - 300000 µg/g body weight), so it is unlikely that females did not respond to treatment because of a dose effect. This is supported by the result that JH-induced increased expression of the conserved gene *krüppel-homolog 1*. One possible explanation for the conflicting results obtained by Schrempf & Heinze 2006 is the increased survival of queen-destined larvae we observed, which may have resulted in changes to caste ratio in the previous experiment. An additional difficulty is that caste proportions obtained from solvent control treatments were not reported in the previous study. Notwithstanding these discrepancies, we are confident, based on our studies of caste phenotypes, which showed extreme differentiation in morphology in early development (Schultner *et al*., 2023), that queen and worker growth and development are highly canalized in this species once caste has been determined. It is possible however that JH-treatment of queens would lead to the increased production of queen-destined eggs, like in other species with maternal caste determination; this remains to be tested. In contrast to females, male development appears to have retained a substantial level of plasticity, perhaps because male morph is determined during larval development. The different degrees of canalization in males and females may also be explained by the evolutionary age of male polymorphism, which though basal to *Cardiocondyla* (Oettler *et al*., 2010), evolved later than queen-worker caste polyphenism.

Juvenile hormone has long been considered the holy grail in ant queen-worker caste polyphenism, even though no studies have followed up on the original reports in *S. invicta*, *P. pallidula* and *M. rubra*. Wheeler herself acknowledged that after this first intense phase of study, investigations into JH have not been as fruitful as initially hoped (Wheeler, 2003). From the studies summarized above as well as our own results, we nevertheless believe that some conclusions can be drawn. First, the mode and the timing of caste determination are clearly critical factors which can influence JH-responsiveness. Second, JH treatment may override even seemingly “hard-wired” caste determination modes such as the hybridogenetic system in *Cataglyphis mauritanica*, but the strength of this effect appears to depend on additional individual- and/or colony-level traits. More studies on species with genetic modes of caste determination are needed to validate these findings. Third, any influence experimental JH manipulation may have on caste and body size can be mediated by mortality rates, as these affect brood:worker ratios and colony size, two important social factors in ant development. To avoid this caveat, experiments should ideally be done with standardized colony sizes. Finally, replication of previous experiments, coupled with modern molecular methods, is needed to evaluate general assumptions about the role of JH in ant caste development, and help move this traditional research field into a new era.

## Methods

### Study species

*Cardiocondyla obscurior* is a myrmicine tramp ant found in the tropics and subtropics. These small ants live in colonies with a few dozen workers and several queens (Heinze, 2017; Oettler, 2020). Queens, workers and males develop via an egg stage and three larval instars, which can be identified by body shape and melanization of the mandibles (Schrempf & Heinze, 2006). Queen and worker-destined eggs and larvae can be distinguished using caste-specific crystalline deposit patterns (Schultner *et al*., 2023). All individuals used in this study derived from laboratory stock colonies of either the Old World population, originally collected in Japan in 2010 (Oyp B), or the New World population, originally collected in Brazil in 2009 (Schrader *et al*., 2014; Errbii *et al*., 2021; Ün *et al*., 2021). Stock and experimental colonies were housed in 9.6×9.6×3 cm plastic boxes with plastered bottoms. In each colony, a plastic insert nest with three chambers and a microscope slide covered with black foil as a lid was placed on a notch in the plaster to prevent the insert nest from slipping and consequently damaging the ants and brood. To hinder workers and emerged brood from escaping, the upper third of the nest was paraffined weekly. Three metal plates were placed inside each colony holding sponges for water supply, honey and insect prey. All colonies were kept in a climate chamber under a 12/12h and 22°C/26°C night/day cycle at 75% relative humidity and fed three times a week with chopped cockroaches (once per week) or fruit flies (twice per week), honey, and water.

#### Hormone manipulations of larvae of unknown caste and sex

In order to replicate the experimental set-up of a previous study (Schrempf & Heinze, 2006), we treated first, second and third instar larvae of unknown sex and caste with 2 µl of either methoprene (1 mg/ml diluted in 70% ethanol) (first instar larvae: n=157, second instar larvae: n=95, third instar larvae: n=40), JH III (1 mg/ml diluted in 70% ethanol) (first instar larvae: n=129, second instar larvae: n=36, third instar larvae: n=24), 70% ethanol (first instar larvae: n=224, second instar larvae: n=102, third instar larvae: n=21), or acetone (first instar larvae: n=66, second instar larvae: n=107, third instar larvae: n=23) using a Hamilton syringe. We also included a handling control (first instar larvae: n=468, second instar larvae: n=223, third instar larvae: n=30). Individuals were treated in groups of five and then transferred on filter paper to rearing nests containing workers from stock colonies. We chose to treat brood in groups because the small size of *C. obscurior* prevents individual treatment of eggs and larvae. For practical reasons, each group received the same amount of 2 µg of JH III or methoprene, which represents doses of ca. 20000 µg/g for L1 larvae, 10000 µg/g for L2 larvae and 1000 µg/g for L3 larvae; these doses are far higher than the effective doses used e.g. by Wheeler & Nijhout 1981 to induced soldier development in Pheidole (50-250 µg/g body weight), but about four times lower than those used by Schrempf & Heinze 2006. Rearing colonies contained between 10-30 workers (∼1:2 worker:brood ratio) and worker numbers were standardized weekly until no more brood remained. The sex and caste/morph of each emerging individual was documented and survival and caste/morph ratios were calculated. Caste/morph ratios were compared between treatments using Fisher’s exact tests in R v4.2.2.

#### Hormone manipulations of queen- and worker-destined late-stage embryos and larvae

To examine whether juvenile hormone can influence caste-specific growth and development after caste has been determined, queen- and worker-destined eggs and larvae (first instar – L1, second instar – L2, third instar – L3) were treated with either juvenile hormone III (JH III) (Sigma-Aldrich, USA), methoprene (Sigma-Aldrich, USA), ethanol (solvent control) or left untreated (handling control). For better identification of caste-specific crystalline patterns in eggs, these were selected in PBT solution (0.3 %). For each developmental stage and caste, individuals were collected from several stock colonies, pooled and then divided into groups of five on filter paper. Each group of five individuals was topically treated with either 1 μl of JH III (1 mg/ml diluted in 70% ethanol), 1 μl of methoprene (1 mg/ml diluted in 70% ethanol), 1 μl of 70% ethanol, or none of the above, using a Hamilton syringe. We reduced doses compared to the previous experiment to minimize mortality and because we found no effect even with higher doses (see next paragraph). This process was conducted twice per treatment day, resulting in 10 individuals being treated per day, treatment, developmental stage and caste. Following treatment, the 10 individuals were transferred on filter paper to a nest containing 15 adult workers collected from several stock colonies the previous day. Two hours after treatment, two individuals were randomly selected from each experimental colony, individually flash-frozen in liquid nitrogen and stored in −80°C for gene expression analyses (see below). The remaining individuals remained in the experimental nests to be reared to adulthood. Treatments were repeated on seven days and a total of 2240 individuals were treated, of which a subset were sampled for gene expression analyses and the rest were allowed to develop until adulthood (n=56 per treatment, developmental stage and caste).

To assess the effect of hormone dose in a separate experiment, we treated groups of five second instar queen- and worker-destined larvae with 2 µl of either methoprene (1 mg/ml diluted in 70% ethanol) (queen-destined: n=10, worker-destined: n=33), JH III (1 mg/ml diluted in 70% ethanol) (queen-destined: n=15, worker-destined: n=15), 70% ethanol (queen-destined: n=11, worker-destined: n=15), or acetone (queen-destined: n=15, worker-destined: n=20) using a Hamilton syringe.

### Caste and morph ratios

Experimental colonies were observed once a week to monitor brood survival and developmental stage. Additionally, the number of workers was counted and any dead or missing workers were replaced from stock colonies. Once the first brood of a colony completed pupation, colonies were checked every other day and when melanization set in, colonies were observed daily to ensure that newly emerged, light-colored adult workers were isolated before they darkened, so as to remain distinguishable from rearing workers. All freshly emerged adults from one experimental colony were transferred together to a smaller round nest with a plastered bottom containing sponges for water supply, a metal plate with honey, and a folded black foil as a shelter, where they were kept for 5-7 days to allow cuticle hardening. The caste and sex of each fully melanized adult was documented and all adults were frozen in Eppendorf tubes at −20 °C for morphometric measurements. From these data, we calculated survival and caste prediction accuracy for each treatment, developmental stage and caste, and compared these using Fisher’s exact tests in R version 4.2.2 (Table 1). Because worker- and male-destined eggs and larvae exhibit similar crystalline deposit patterns (pers. obs. E. Schultner), in each treatment a small proportion of individuals developed into males (31 males in total, Table 1). These were excluded prior to survival and caste prediction accuracy analyses. Note that the survival and caste prediction accuracy results from the handling control have been published as supplemental data in a previous study (Schultner *et al*., 2023); we nevertheless include them here for comparison.

### Development time

Development time was defined as the time in days between treatment and emergence of the first adult from each experimental colony. Development time was calculated separately for each treatment, developmental stage and caste. In cases where more than one caste emerged from an experimental colony, development time was calculated for the first hatched individual of each caste. In late-stage embryos, L1 and L2 larvae, development time was compared between castes and treatments using a linear regression with caste and treatment as explanatory variables, as well as an interaction term to account for caste-specific responses to treatment: *development time ∼ caste * treatment*. Model fit was assessed using residual tests implemented in the DHarma package in R v4.2.2 (Hartig, 2020). Tukey-corrected pairwise p-values were calculated using the emmeans package in R v4.2.2 (Lenth *et al*., 2020). In L3 larvae, Kruskal Wallis tests (function kruskal.test in R v4.2.2) were used to test the effect of caste and treatment on development time separately, as data did not follow a normal distribution.

### Body size measurements

To assess the effect of hormone treatment on adult body size, all adults which emerged in experimental colonies were dried under a stereomicroscope light and pinned for subsequent measurements (Table 1). Pinned individuals were photographed at 200x using a Keyence stereomicroscope connected to a camera (Keyence VHX). Head width (HW), head length (HL), mesosoma width (MW), mesosoma length (ML), and petiole width (PW) were measured from photographs after appropriate scaling using ImageJ (v. 1.53 s). In the cases in which adult female caste did not align with presumed caste based on crystalline deposit patterns (queens: 12.8% (51/397) of individuals, workers: 3% (10/331) of individuals, Table 1), the emerging adults were nonetheless measured and included in subsequent analyses. Similarly, all emerged males were measured and included in analyses. The final data set contained 759 individuals (Queens: late-stage embryos - handling control: n=9, solvent control: n=13, JH III: n=8, methoprene: n=13; L1 larvae - handling control: n=14, solvent control: n=23, JH III: n=18, methoprene: n=15; L2 larvae - handling control: n=26, solvent control: n=25, JH III: n=28, methoprene: n=25; L3 larvae - handling control: n=43, solvent control: n=38, JH III: n=27, methoprene: n=31; Workers: late-stage embryos - handling control: n=27, solvent control: n=16, JH III: n=20, methoprene: n=20; L1 larvae – handling control: n=24, solvent control: n=19, JH III: n=19, methoprene: n=18; L2 larvae - handling control: n=29, solvent control: n=27, JH III: n=27, methoprene: n=29; L3 larvae - handling control: n=23, solvent control: n=24, JH III: n=29, methoprene: n=21; Ergatoid males: late-stage embryos - handling control: n=1, solvent control: n=1, JH III: n=4; L1 larvae - handling control: n=1, solvent control: n=2, JH III: n=1, methoprene: n=3; L2 larvae - handling control: n=1, solvent control: n=2, JH III: n=1, methoprene: n=1; L3 larvae - handling control: n=2, solvent control: n=5, JH III: n=1, methoprene: n=5). A principal component analysis (PCA) of the five measured traits was done using the NIPALS algorithm implemented in the pcaMethods package in R v4.2.2 (Stacklies *et al*., 2007) to estimate missing values. Principal component 1 scores were compared between the three castes (queens, workers, males), the four developmental stages (late-stage embryos, L1, L2, L3 larvae) and between treatments using a linear regression, with caste, treatment and developmental stage as explanatory variables and an interaction term to account for caste-specific responses to treatment: *PC score ∼ caste * treatment + developmental stage*. Residual tests implemented in the DHarma package in R v4.2.2 (Hartig, 2020) were used to assess model fit. There was no significant effect of developmental stage on adult body size (Table 2), so this term was removed from the model before pairwise p-values were calculated and Tukey-corrected for multiple comparisons using the emmeans package in R v4.2.2 (Lenth *et al*., 2020).

Since the treatment was applied in a single topical application in each developmental stage, it is possible that an effect on body size was diluted over time. To assess the short-term effect of hormonal manipulation, we repeated hormone treatments using only L2 larvae, which were measured once they reached the L3 stage. As in the previous experiment, L2 larvae of each caste were chosen based on crystalline deposit patterns and treated in groups of five with 1 µl of either JH III, methoprene or ethanol as a solvent control, or subjected to a handling control. The five larvae were transferred to a nest with 12 adult workers originating from several stock colonies which had been set up one day earlier. This procedure was repeated seven times, for a total of 35 L2 larvae per treatment and caste. After treatment, nests were checked every day for moulted larvae and all larvae that had reached the L3 stage were anaesthetized under CO_2_ for 4h. Larvae were then placed on their backs and photographed at 200x using a Keyence stereomicroscope connected to a camera (Keyence VHX). The order in which the pictures were taken was randomized to buffer the effect of time as larvae were more likely to die, lose water (change of body volume) or wake up after a long waiting period. Larval head capsule width, body width and body length were measured after appropriate scaling using ImageJ (v. 1.53 t). As some larvae died or were damaged in the pre-imaging process, the final data set consisted of 29-35 individuals per caste and treatment (Queens - handling control: n= 32, solvent control: n=29, JH III: n=35, methoprene: n=32; Workers - handling control: n=32, solvent control: n=32, JH III: n=30, methoprene: n=28). Like in adults, a principal component analysis including all traits was run using the NIPALS algorithm implemented in the pcaMethods package in R v4.2.2 (Stacklies *et al*., 2007) to estimate missing values. Principal component 1 scores were then compared between castes and between treatments using a linear regression, with caste and treatment as explanatory variables and an interaction term to account for caste-specific responses to treatment: *PC 1 score ∼ caste * treatment*. Residual tests implemented in the DHarma package in R v4.2.2 (Hartig, 2020) were used to test model fit. Pairwise p-values were calculated and Tukey-corrected for multiple comparisons using the emmeans package in R v4.2.2 (Lenth *et al*., 2020).

#### Hormone manipulations of early-stage embryos

To test whether caste/morph determination and differentiation can be manipulated early in development before caste-specific crystalline deposits are visible, we treated female and male eggs which were at most 24 hours old. To obtain female eggs, experimental colonies containing three mated queens of unknown age, 10 adult workers and five worker pupae were set up from stock colonies (n=25 colonies). To obtain male eggs, experimental colonies containing three queen pupae prior to mating, 15 adult workers and five worker pupae were set up from stock colonies (n=40 colonies). Ants exhibit haplodiploid sex determination, so unmated queens only produce male-destined eggs. Experimental colonies were monitored once per week until the first eggs were observed. Once experimental colonies began producing sufficient numbers of eggs, eggs were removed daily to ensure that only eggs which were at most 24 hours old were included in experiments. Eggs from several experimental colonies were pooled and then divided into groups of five on filter paper, after which they were topically treated with either 1 μl of methoprene (1 mg/ml diluted in 70% ethanol) or 1 μl of 70% ethanol using a Hamilton syringe. This was done in parallel for both sexes to avoid bias. Treated eggs were transferred on filter paper to rearing colonies containing 15 workers randomly collected from stock colonies; rearing colonies were always set up one day before egg transfer to allow workers to acclimate. Each rearing colony contained 5-25 eggs, as the number of treated eggs depended on the fecundity of queens, which is much higher in mated compared to unmated queens and varies over time (pers. obs. E. Schultner). In total, we treated 650 early-stage male embryos (solvent control: n=325, methoprene: n=325) and 570 early-stage female embryos (solvent control: n=285, methoprene: n=285). Rearing colonies were monitored twice per week and fed in the same manner as stock colonies. Freshly emerged adults were transferred to a smaller nest with a plastered bottom and sponges for water supply, a metal plate with honey, and a folded black foil as a shelter, and kept for 3-5 days to allow cuticle hardening. The caste/morph of each fully melanized adult was documented and all adults were frozen in Eppendorf tubes at −20 °C for morphometric measurements. From these data, survival and caste/morph ratios were calculated and survival and treatment effects were analysed using Fisher’s exact tests in R v4.2.2.

### Body size measurements

Body size measurements of adults emerging from treatment of early-stage embryos were obtained as described above for later developmental stages. The final data set consisted of 74 individuals (queens - solvent control: n=1, methoprene: n=4; workers - solvent control: n=24, methoprene: n=24; ergatoid males - solvent control: n=10, methoprene: n=11). A principal component analysis using the NIPALS algorithm to estimate missing values implemented in the pcaMethods package in R v4.2.2 (Stacklies *et al*., 2007) was run on all traits. Principal component 1 and 2 scores were compared between castes and treatments using a linear regression, with caste and treatment as explanatory variables, as well as an interaction term to account for caste-specific responses to treatment: *PC 1/2 score ∼ caste * treatment.* Residual tests implemented in the DHarma package in R v4.2.2 (Hartig, 2020) were used to test model fit and pairwise p-values were calculated and Tukey-corrected for multiple comparisons using the emmeans package in R v4.2.2 (Lenth *et al*., 2020).

#### Gene expression analysis

qPCR was used to assess the effect of sex, caste and treatment on the expression of the JH-response gene *Krüppel-homolog 1* (*kr-h1*). *C. obscurior* sequences for *kr-h1* were retrieved by running a BLAST search of *kr-h1* protein sequences obtained for *Drosophila melanogaster* (CG45074, www.flybase.org) and *Apis mellifera* (GB45427, www.hymenopteragenome.org) against the *C. obscurior* 1.4 genome (Schrader et al. 2014). Intron-spanning primers for the extracted *C. obscurior* ortholog (Cobs_15554) were designed in Geneious Prime vR10 (https://www.geneious.com). Specific primers (*kr-h1* forward primer sequence: CTTGGTGTGCAGCCCGGACC, kr-h1 reverse primer sequence: ACCGGTACGGATCCTCGCCC) were tested on pooled samples of queen, worker, ergatoid male and winged male DNA in a temperature gradient PCR (60°C, 63°C, 66°C) and PCR products subsequently sequenced to confirm amplification of the desired transcript. Primer efficiency was tested in a five-step dilution series and was confirmed to be within the acceptable range.

To establish a timeline of *kr-h1* expression and to investigate sex-specific responses to JH, second instar male and female larvae (of unknown caste) were treated in groups of five with 2 µl of either JH III (1 mg/ml diluted in 70% ethanol) or 70% ethanol using a Hamilton syringe. We further included a handling control, which consisted of larvae which were removed from colonies and immediately flash-frozen in liquid nitrogen and stored at −80°C. Female larvae were collected from stock colonies containing mated queens whereas male larvae were collected from colonies containing only unmated queens. After treatment, larvae were moved to rearing nests containing workers from stock colonies, which had been set up the day before. One, two, six and 24 hours after treatment, treated larvae were removed from rearing colonies, individually flash-frozen in liquid nitrogen and stored in −80°C. We ensured that only larvae which had been accepted by workers, i.e. which had been moved inside the nest after treatment, were included. The final sample consisted of 177 females (handling control - 0h: n=19; ethanol - 1h: n=14, 2h: n=11, 6h: n=13, 24h: n=10; JH III - 1h: n=12, 2h: n=12, 6h: n=15, 24h: n=15; methoprene – 1h: n=16, 2h: n=9, 6h: n=13, 24h: n=18) and 111 males (handling control - 0h: n=21; ethanol - 1h: n=13, 2h: n=13, 6h: n=8, 24h: n=6; JH III – 1h: n=13, 2h: n=12, 6h: n=6, 24h: n=19).

To assess caste-specific responses to hormone treatment, *kr-h1* expression was measured in queen- and worker-destined late-stage embryos, first instar, second instar and third instar larvae. All samples were collected from the experiment described in the above section “Hormone manipulations of queen- and worker-destined late-stage embryos and larvae”, two hours after treatment. This timepoint was chosen based on the timeline of *kr-h1* expression in female embryos of unknown caste described above (Figure 4). The final data set consisted of 5-14 individuals per treatment, developmental stage and caste (Queens: late-stage embryos - handling control: n=14, solvent control: n=9, JH: n=12, methoprene: n=10; L1 larvae - handling control: n=5, solvent control: n=8, JH: n=10, methoprene: n=10; L2 larvae - handling control: n=5, solvent control: n=8, JH: n=8, methoprene: n=8; L3 larvae - handling control: n=7, solvent control: n=9, JH: n=9, methoprene: n=9; Workers: late-stage embryos - handling control: n=12, solvent control: n=10, JH: n=9, methoprene: n=9; L1 larvae - handling control: n=6, solvent control: n=7, JH: n=7, methoprene: n=8; L2 larvae - handling control: n=5, solvent control: n=6, JH: n=8, methoprene: n=6; L3 larvae - handling control: n=6, solvent control: n=9, JH: n=7, methoprene: n=5).

Each sample was homogenized with ceramic beads in a shaker and total RNA was extracted using a modified protocol of the ReliaPrep^TM^ RNA Cell Miniprep System (Promega, USA) and cDNA was synthesized with the iScript^TM^ gDNA Clear cDNA Synthesis Kit (Bio-Rad, USA), following the manufacturer’s instruction. cDNA concentration was measured using the Qubit dsDNA HS assay kit (ThermoFisher Scientific, USA) and standardized to 1.5 - 3 ng/µl, depending on developmental stage. Each qPCR reaction was run in 10 µl (5 µl SYBR green, 3 µl purified H_2_O, 0.5 µl forward primer, 0.5 ml reverse primer, 1 µl cDNA) in a qPCR machine (CFX, Bio-Rad, USA). Three technical replicates were run for each sample and gene. In addition to kr-h1 and the housekeeping gene y45, in all samples we analyzed expression of the two sex-specific *doublesex* isoforms *dsx^f^* and *dsx^m^,* which vary in their expression between males and females (Klein *et al*., 2016). This was done to identify males in samples collected from stock colonies containing mated queens, as males cannot be distinguished morphologically from workers before the pupal stage. All male samples thus identified were excluded before statistical analyses. For amplification of *dsx^F^, dsx^M^* and the housekeeper *y45*, primer sequences designed for a previous study were used (Klein et al. 2016).The housekeeping gene *y45* was used for normalization according to the deltaCq method (Schmittgen & Livak, 2008). For timeline data, differences in *kr-h1* expression between treatments were compared separately for each sex and timepoint with linear regressions, using log-transformed expression as a response and treatment as explanatory variable: kr-h1 expression_sex,timepoint_ ∼ treatment. For caste-specific responses to treatment *kr-h1* expression was compared separately for each developmental stage, using using log-transformed expression as a response and caste and treatment as explanatory variables: kr-h1 expression_developmental stage_ ∼ caste + treatment. For all analyses, model fit was assessed using residual tests implemented in the DHarma package in R v4.2.2 (Hartig, 2020) and pairwise p-values were calculated and Tukey-corrected for multiple comparisons using the emmeans package in R v4.2.2 (Lenth *et al*., 2020).

## Acknowledgements

The authors thank H. Lowack, B. Grüneberg, S. Stewart, J. Wallner, L. Drechsel, H. Sammereier & G. Gebhard for help with experiments. This work was funded by a research stipend from the German Academic Exchange Service DAAD to J.B., a research grant from the German Research Foundation DFG to J.O. (Oe549/3-1,2), and a research stipend from the Bavarian State Ministry of Science and Art to E.S. Funders had no role in the design of the study, data collection or interpretation of results.

