## Supplement Figures for "Sex- and caste-specific developmental responses to juvenile hormone in an ant with maternal caste determination"

Figure S1: Boxplots comparing individual traits measured in L3 larvae emerging from treatments of L2 larvae

a) Head width
b) Body width
c) Body length

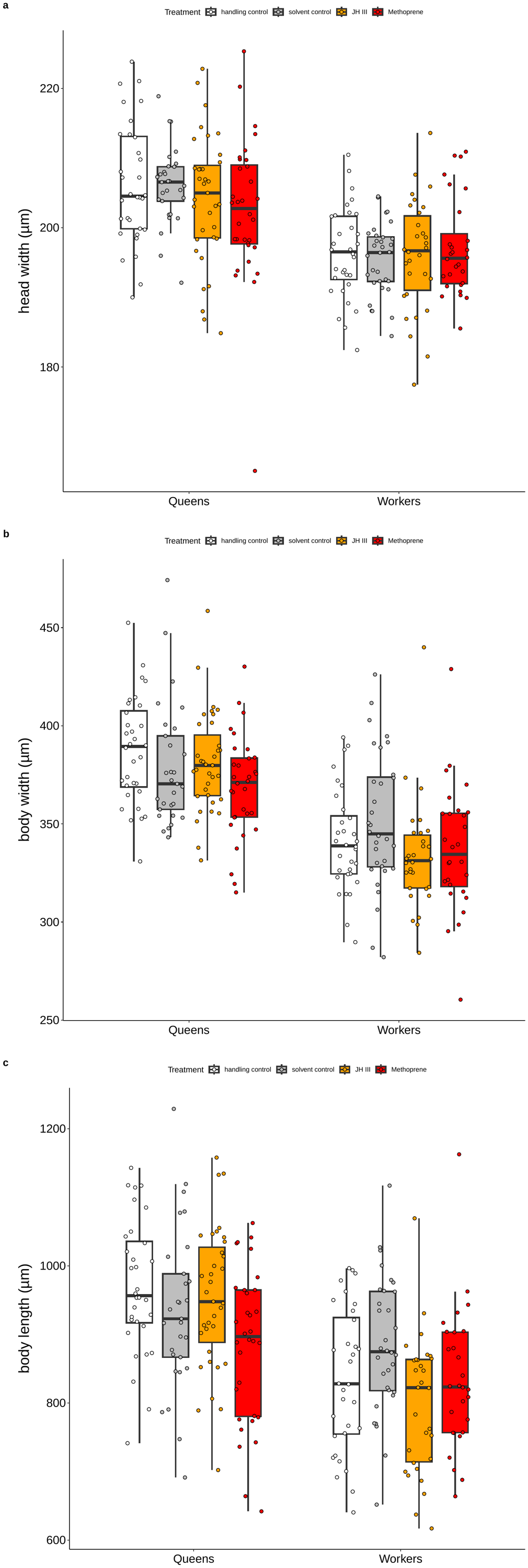

Figure S2: Boxplots comparing individual traits measured in adult queens, workers and males emerging from hormone treatment of late-stage embryos and larvae

a) Head width, b) head length, c) thorax width, d) thorax length, e) petiole width

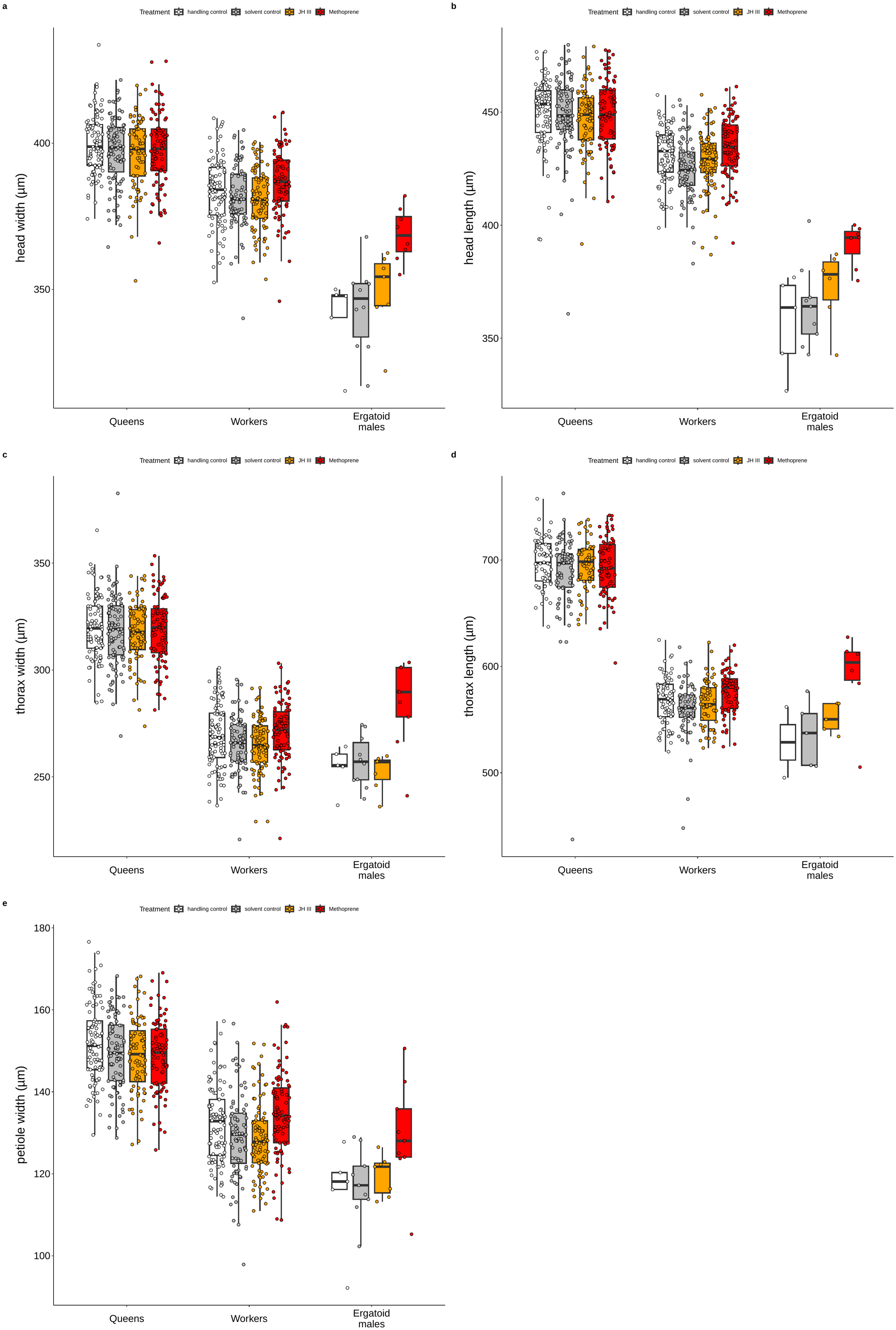

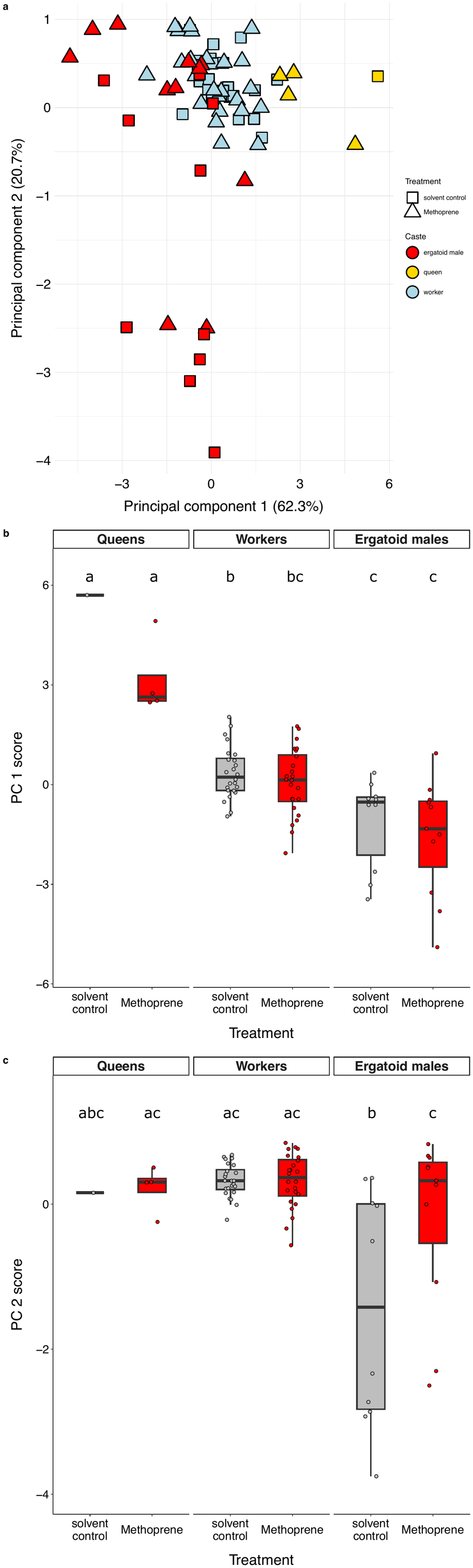
Figure S4: Body size of adult queens, workers and males after hormone treatment of early-stage embryos

a) Principal component analysis separates individuals by overall body size on PC1 and by petiole width on PC2. See table S7 for loadings.

b) PC 1 scores plotted by caste and treatment show that queens are the largest caste, and workers are larger than males. Overall size is not affected by hormone treatment in any of the three groups. See supplement table S6 for Tukey-corrected pairwise p-values.

c) PC 2 scores plotted by caste and treatment show a significant difference between treatment in males but not females.

Figure S5: Boxplots comparing individual traits measured in adult queens, workers and males emerging from hormone treatment of early-stage embryos

a) Head width, b) head length, c) thorax width, d) thorax length, e) petiole width

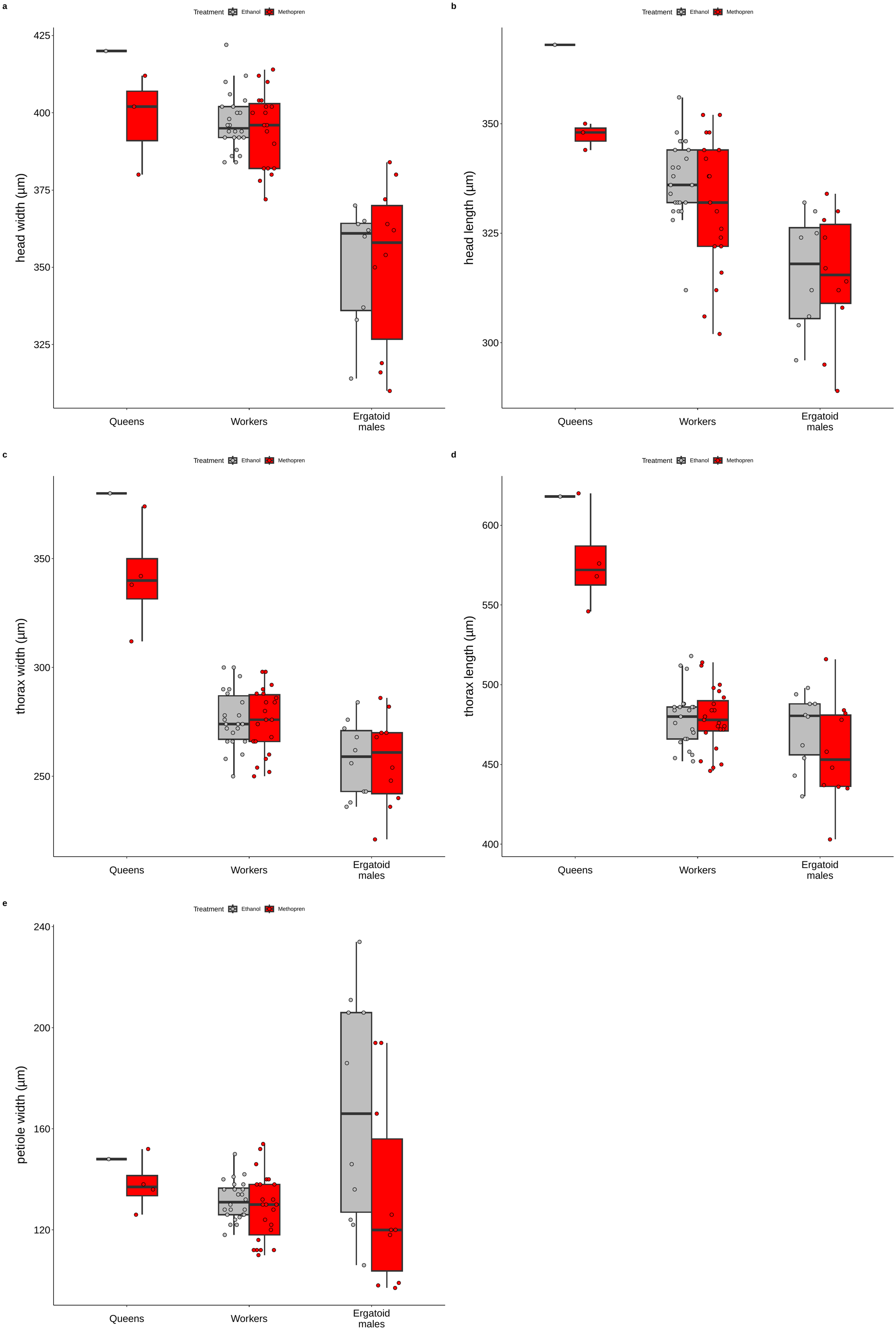

Figure S6: Normalized expression of *krüppel-homolog 1* in queen- and worker-destined late-stage embryos and larvae after hormone treatment

(Ctrl=handling control, EtOH=solvent control, JH=JH III, Met=Methoprene)

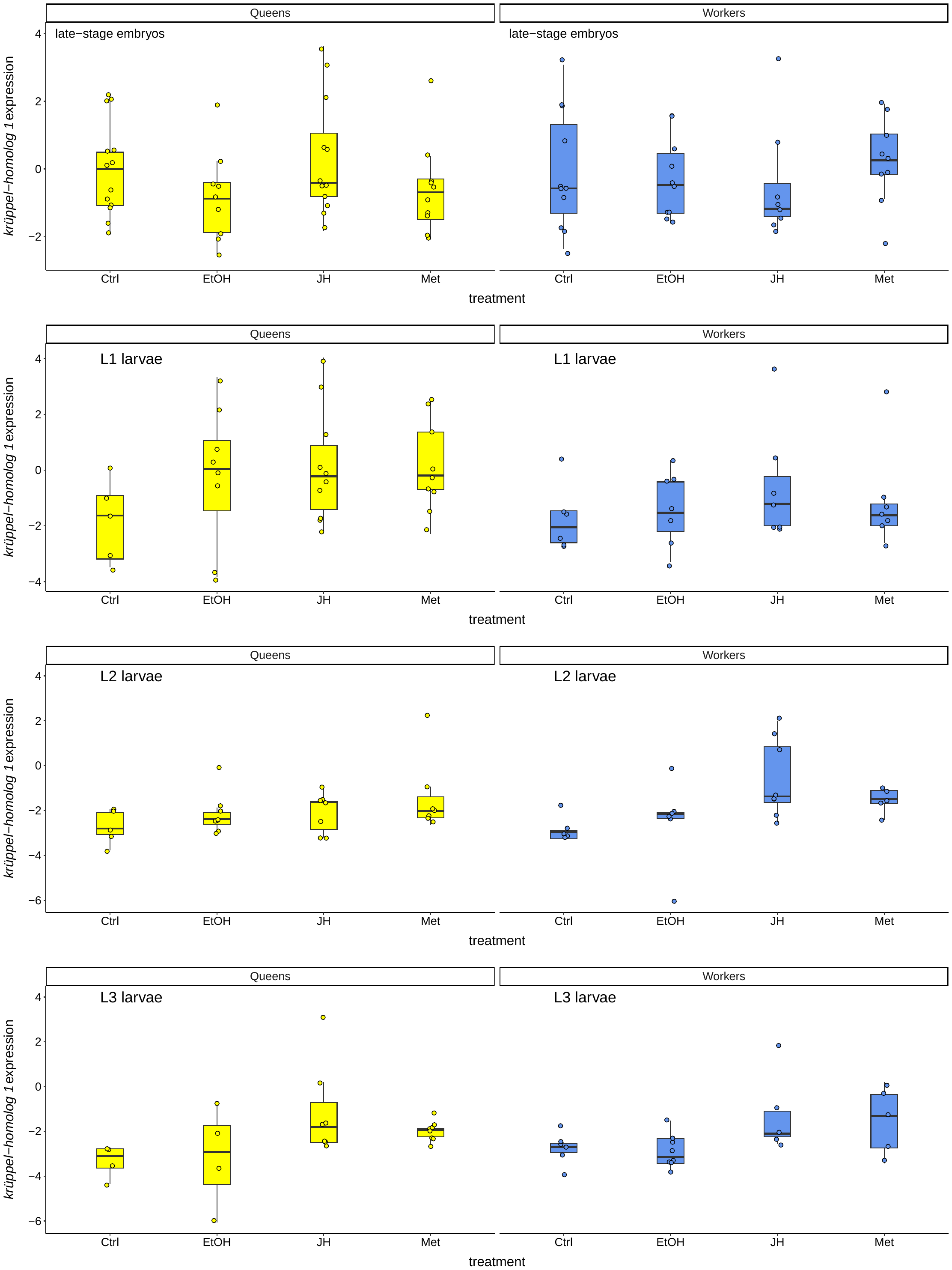
