## Supplement Tables for "Sex- and caste-specific developmental responses to juvenile hormone in an ant with maternal caste determination"

| **Developmental stage** | **Caste** | **Treatment** | **n** | **Survival** | **Queens** | **Workers** | **Ergatoid Males** | **Winged Males** | **Proportion winged individuals** | **Fisher's test within stages & between treatments (p<0.05)** |
| --- | --- | --- | --- | --- | --- | --- | --- | --- | --- | --- |
| 1st instar larva | unknown | handling control | 123 | 34.1% (42/123) | 20 | 22 | 0 | 0 | 47.6% (20/42) | ab |
|  |  | solvent control (70% ethanol) | 224 | 9.8% (22/224) | 9 | 13 | 0 | 0 | 40.9% (9/22) | ab |
|  |  | solvent control (acetone) | 66 | 31.8% (21/66) | 10 | 11 | 0 | 0 | 47.6% (10/21) | ab |
|  |  | JH III (in 70% ethanol) | 129 | 11.6% (15/129) | 3 | 12 | 0 | 0 | 20% (3/15) | a |
|  |  | Methoprene (in 70% ethanol) | 157 | 11.4% (18/157) | 12 | 5 | 0 | 1 | 72.2% (13/18) | b |
| 2nd instar larva | unknown | handling control | 113 | 49.6% (56/113) | 10 | 43 | 3 | 0 | 17.9% (10/56) | a |
|  |  | solvent control (70% ethanol) | 102 | 47.1% (48/102) | 13 | 33 | 2 | 0 | 27.1% (13/48) | a |
|  |  | solvent control (acetone) | 107 | 46.7% (50/107) | 1 | 44 | 5 | 0 | 2% (1/50) | b |
|  |  | JH III (in 70% ethanol) | 36 | 50% (18/36) | 4 | 14 | 0 | 0 | 22.2% (4/18) | a |
|  |  | Methoprene (in 70% ethanol) | 95 | 51.6% (49/95) | 9 | 39 | 1 | 0 | 18.4% (9/36) | a |
| 3rd instar larva | unknown | handling control | 30 | 83.3% (25/30) | 6 | 18 | 1 | 0 | 24% (6/25) | a |
|  |  | solvent control (70% ethanol) | 21 | 90.5% (19/21) | 4 | 15 | 0 | 0 | 21.1% (4/19) | a |
|  |  | solvent control (acetone) | 23 | 82.6% (19/23) | 4 | 15 | 0 | 0 | 21.1% (4/19) | a |
|  |  | JH III (in 70% ethanol) | 24 | 75% (18/24) | 4 | 11 | 1 | 2 | 33.3% (6/18) | a |
|  |  | Methoprene (in 70% ethanol) | 40 | 40% (16/40) | 7 | 9 | 0 | 0 | 43.8% (7/16) | a |
| 2nd instar larvae | queen | solvent control (70% ethanol) | 11 | 81.1% (9/11) | 9 | 0 | 0 | 0 | 100% (9/9) | a |
|  |  | solvent control (acetone) | 15 | 73.3% (11/15) | 11 | 0 | 0 | 0 | 100% (11/11) | a |
|  |  | JH III (in 70% ethanol) | 15 | 66.7% (10/15) | 10 | 0 | 0 | 0 | 100% (10/10) | a |
|  |  | Methoprene (in 70% ethanol) | 10 | 70% (7/10) | 7 | 0 | 0 | 0 | 100% (7/7) | a |
|  | worker | solvent control (70% ethanol) | 15 | 46.7% (7/15) | 0 | 7 | 0 | 0 | 0% (7/7) | a |
|  |  | solvent control (acetone) | 20 | 60% (12/20) | 0 | 11 | 1 | 0 | 0% (12/12) | a |
|  |  | JH III (in 70% ethanol) | 15 | 60% (9/15) | 0 | 9 | 0 | 0 | 0% (9/9) | a |
|  |  | Methoprene (in 70% ethanol) | 33 | 72.7% (24/33) | 0 | 21 | 3 | 0 | 0% (24/24) | a |

**Table S1:** Survival and caste/morph ratios following treatment of larvae with 2 µl of 1 mg/ml hormone solution

| contrast | estimate | SE | df | t.ratio | p.value |
| --- | --- | --- | --- | --- | --- |
| QU Ctrl - WO Ctrl | 2.0591 | 0.2856 | 242 | 7.2111 | <0.001 |
| QU Ctrl - QU EtOH | 0.3542 | 0.2928 | 242 | 1.2097 | 0.9285 |
| QU Ctrl - WO EtOH | 1.6589 | 0.2856 | 242 | 5.8095 | <0.001 |
| QU Ctrl - QU JH | 0.3112 | 0.2794 | 242 | 1.1141 | 0.9533 |
| QU Ctrl - WO JH | 2.4156 | 0.2903 | 242 | 8.3218 | <0.001 |
| QU Ctrl - QU Metho | 0.9806 | 0.2856 | 242 | 3.4341 | 0.0158 |
| QU Ctrl - WO Metho | 2.1010 | 0.2956 | 242 | 7.1082 | <0.001 |
| WO Ctrl - QU EtOH | -1.7049 | 0.2928 | 242 | -5.8218 | <0.001 |
| WO Ctrl - WO EtOH | -0.4002 | 0.2856 | 242 | -1.4016 | 0.8559 |
| WO Ctrl - QU JH | -1.7479 | 0.2794 | 242 | -6.2567 | <0.001 |
| WO Ctrl - WO JH | 0.3564 | 0.2903 | 242 | 1.2280 | 0.9229 |
| WO Ctrl - QU Metho | -1.0785 | 0.2856 | 242 | -3.7770 | 0.0049 |
| WO Ctrl - WO Metho | 0.0419 | 0.2956 | 242 | 0.1417 | 1.0000 |
| QU EtOH - WO EtOH | 1.3047 | 0.2928 | 242 | 4.4552 | <0.001 |
| QU EtOH - QU JH | -0.0430 | 0.2868 | 242 | -0.1500 | 1.0000 |
| QU EtOH - WO JH | 2.0613 | 0.2974 | 242 | 6.9300 | <0.001 |
| QU EtOH - QU Metho | 0.6264 | 0.2928 | 242 | 2.1389 | 0.3931 |
| QU EtOH - WO Metho | 1.7468 | 0.3026 | 242 | 5.7720 | <0.001 |
| WO EtOH - QU JH | -1.3477 | 0.2794 | 242 | -4.8241 | <0.001 |
| WO EtOH - WO JH | 0.7567 | 0.2903 | 242 | 2.6067 | 0.1586 |
| WO EtOH - QU Metho | -0.6783 | 0.2856 | 242 | -2.3754 | 0.2582 |
| WO EtOH - WO Metho | 0.4421 | 0.2956 | 242 | 1.4957 | 0.8092 |
| QU JH - WO JH | 2.1043 | 0.2842 | 242 | 7.4048 | <0.001 |
| QU JH - QU Metho | 0.6694 | 0.2794 | 242 | 2.3961 | 0.2480 |
| QU JH - WO Metho | 1.7898 | 0.2896 | 242 | 6.1801 | <0.001 |
| WO JH - QU Metho | -1.4350 | 0.2903 | 242 | -4.9435 | <0.001 |
| WO JH - WO Metho | -0.3146 | 0.3001 | 242 | -1.0481 | 0.9664 |
| QU Metho - WO Metho | 1.1204 | 0.2956 | 242 | 3.7906 | 0.0046 |

**Table S2:** Tukey-corrected pairwise p-values comparing body size of L3 larvae emerging from treatments of L2 larvae

Morphometric measurements were subjected to a principal component analysis followed by linear regression of principal component 1 scores. (QU=queen, WO=worker, Ctrl=handling control, EtOH=solvent control, JH=juvenile hormone III, Metho=Methoprene)

**Table S3:** Loadings of size measurements of L3 larvae emerging from treatments of L2 larvae on principal components 1-3

| Trait | PC1 | PC2 | PC3 |
| --- | --- | --- | --- |
| Head width | 0.440498 | 0.8770605 | 0.1916411 |
| Body length | 0.612554 | -0.4496885 | 0.6500445 |
| Body width | 0.6563072 | -0.1689528 | -0.7353338 |

**Table S4:** Tukey-corrected pairwise p-values comparing body size of adults emerging from treatments of late-stage embryos and larvae

Morphometric measurements were subjected to a principal component analysis followed by linear regression of principal component 1 scores. (EM=ergatoid male, QU=queen, WO=worker, Ctrl=handling control, EtOH=solvent control, JH=juvenile hormone III, Metho=Methoprene)

| contrast | estimate | SE | df | t.ratio | p.value |
| --- | --- | --- | --- | --- | --- |
| EM Ctrl - QU Ctrl | -6.4229528 | 0.51077875 | 747 | -12.574824 | <0.001 |
| EM Ctrl - WO Ctrl | -3.6245119 | 0.5093709 | 747 | -7.1156634 | <0.001 |
| EM Ctrl - EM EtOH | -0.5853907 | 0.60923732 | 747 | -0.9608583 | 0.99838874 |
| EM Ctrl - QU EtOH | -6.2072896 | 0.50984708 | 747 | -12.174807 | <0.001 |
| EM Ctrl - WO EtOH | -3.2420991 | 0.51169637 | 747 | -6.3359822 | <0.001 |
| EM Ctrl - EM JH | -0.8065053 | 0.65130209 | 747 | -1.2382969 | 0.98571987 |
| EM Ctrl - QU JH | -6.0671436 | 0.51256339 | 747 | -11.836865 | <0.001 |
| EM Ctrl - WO JH | -3.3092305 | 0.51036286 | 747 | -6.4840739 | <0.001 |
| EM Ctrl - EM Metho | -2.8591987 | 0.62041692 | 747 | -4.6085118 | <0.001 |
| EM Ctrl - QU Metho | -6.2514611 | 0.51203097 | 747 | -12.209147 | <0.001 |
| EM Ctrl - WO Metho | -3.8781541 | 0.51137678 | 747 | -7.5837508 | <0.001 |
| QU Ctrl - WO Ctrl | 2.79844092 | 0.15956243 | 747 | 17.5382192 | <0.001 |
| QU Ctrl - EM EtOH | 5.83756211 | 0.37036679 | 747 | 15.7615699 | <0.001 |
| QU Ctrl - QU EtOH | 0.21566319 | 0.16107608 | 747 | 1.33889024 | 0.97376094 |
| QU Ctrl - WO EtOH | 3.1808537 | 0.16683714 | 747 | 19.0656213 | <0.001 |
| QU Ctrl - EM JH | 5.61644749 | 0.43611451 | 747 | 12.8783781 | <0.001 |
| QU Ctrl - QU JH | 0.35580921 | 0.16947768 | 747 | 2.0994459 | 0.62337139 |
| QU Ctrl - WO JH | 3.1137223 | 0.16270128 | 747 | 19.1376638 | <0.001 |
| QU Ctrl - EM Metho | 3.56375416 | 0.38848244 | 747 | 9.17352712 | <0.001 |
| QU Ctrl - QU Metho | 0.17149169 | 0.16786055 | 747 | 1.02163185 | 0.99720053 |
| QU Ctrl - WO Metho | 2.54479873 | 0.16585435 | 747 | 15.3435752 | <0.001 |
| WO Ctrl - EM EtOH | 3.03912119 | 0.36842278 | 747 | 8.2490045 | <0.001 |
| WO Ctrl - QU EtOH | -2.5827777 | 0.15655442 | 747 | -16.497635 | <0.001 |
| WO Ctrl - WO EtOH | 0.38241278 | 0.16247589 | 747 | 2.35365858 | 0.43935254 |
| WO Ctrl - EM JH | 2.81800658 | 0.43446478 | 747 | 6.4861565 | <0.001 |
| WO Ctrl - QU JH | -2.4426317 | 0.16518616 | 747 | -14.787145 | <0.001 |
| WO Ctrl - WO JH | 0.31528138 | 0.15822607 | 747 | 1.99260069 | 0.69850709 |
| WO Ctrl - EM Metho | 0.76531324 | 0.38662952 | 747 | 1.97944853 | 0.70742023 |
| WO Ctrl - QU Metho | -2.6269492 | 0.1635266 | 747 | -16.064354 | <0.001 |
| WO Ctrl - WO Metho | -0.2536422 | 0.16146656 | 747 | -1.5708651 | 0.91909971 |
| EM EtOH - QU EtOH | -5.6218989 | 0.36908086 | 747 | -15.232161 | <0.001 |
| EM EtOH - WO EtOH | -2.6567084 | 0.37163128 | 747 | -7.148775 | <0.001 |
| EM EtOH - EM JH | -0.2211146 | 0.54815239 | 747 | -0.4033817 | 0.99999975 |
| EM EtOH - QU JH | -5.4817529 | 0.37282417 | 747 | -14.70332 | <0.001 |
| EM EtOH - WO JH | -2.7238398 | 0.36979303 | 747 | -7.3658496 | <0.001 |
| EM EtOH - EM Metho | -2.273808 | 0.51107121 | 747 | -4.449102 | <0.001 |
| EM EtOH - QU Metho | -5.6660704 | 0.37209185 | 747 | -15.227612 | <0.001 |
| EM EtOH - WO Metho | -3.2927634 | 0.37119111 | 747 | -8.8708034 | <0.001 |
| QU EtOH - WO EtOH | 2.96519051 | 0.16396264 | 747 | 18.0845493 | <0.001 |
| QU EtOH - EM JH | 5.40078431 | 0.43502297 | 747 | 12.4149407 | <0.001 |
| QU EtOH - QU JH | 0.14014602 | 0.16664873 | 747 | 0.84096663 | 0.99953574 |
| QU EtOH - WO JH | 2.89805911 | 0.15975238 | 747 | 18.1409449 | <0.001 |
| QU EtOH - EM Metho | 3.34809097 | 0.38725666 | 747 | 8.64566402 | <0.001 |
| QU EtOH - QU Metho | -0.0441715 | 0.16500389 | 747 | -0.2676998 | 1 |
| QU EtOH - WO Metho | 2.32913554 | 0.16296252 | 747 | 14.2924612 | <0.001 |
| WO EtOH - EM JH | 2.4355938 | 0.43718887 | 747 | 5.57103339 | <0.001 |
| WO EtOH - QU JH | -2.8250445 | 0.17222348 | 747 | -16.403364 | <0.001 |
| WO EtOH - WO JH | -0.0671314 | 0.16555951 | 747 | -0.405482 | 0.99999973 |
| WO EtOH - EM Metho | 0.38290046 | 0.38968814 | 747 | 0.98258176 | 0.99802612 |
| WO EtOH - QU Metho | -3.009362 | 0.17063239 | 747 | -17.636523 | <0.001 |
| WO EtOH - WO Metho | -0.636055 | 0.16865916 | 747 | -3.7712447 | 0.0095204 |
| EM JH - QU JH | -5.2606383 | 0.43820333 | 747 | -12.005017 | <0.001 |
| EM JH - WO JH | -2.5027252 | 0.43562734 | 747 | -5.7451058 | <0.001 |
| EM JH - EM Metho | -2.0526933 | 0.56055159 | 747 | -3.6619169 | 0.01412323 |
| EM JH - QU Metho | -5.4449558 | 0.43758044 | 747 | -12.443325 | <0.001 |
| EM JH - WO Metho | -3.0716488 | 0.43681477 | 747 | -7.0319251 | <0.001 |
| QU JH - WO JH | 2.75791309 | 0.1682201 | 747 | 16.3946702 | <0.001 |
| QU JH - EM Metho | 3.20794495 | 0.39082592 | 747 | 8.20811716 | <0.001 |
| QU JH - QU Metho | -0.1843175 | 0.17321508 | 747 | -1.0640963 | 0.99599233 |
| QU JH - WO Metho | 2.18898952 | 0.17127161 | 747 | 12.7808079 | <0.001 |
| WO JH - EM Metho | 0.45003186 | 0.38793546 | 747 | 1.16006889 | 0.99162384 |
| WO JH - QU Metho | -2.9422306 | 0.16659077 | 747 | -17.661426 | <0.001 |
| WO JH - WO Metho | -0.5689236 | 0.16456909 | 747 | -3.4570499 | 0.02841675 |
| EM Metho - QU Metho | -3.3922625 | 0.39012739 | 747 | -8.6952686 | <0.001 |
| EM Metho - WO Metho | -1.0189554 | 0.3892684 | 747 | -2.6176166 | 0.27156747 |
| QU Metho - WO Metho | 2.37330704 | 0.16967158 | 747 | 13.9876519 | <0.001 |

**Table S5:** Loadings of size measurements of adults emerging from treatments of late-stage embryos and larvae on principal components 1-5

| Trait | PC1 | PC2 | PC3 | PC4 | PC5 |
| --- | --- | --- | --- | --- | --- |
| Head width | 0.4642101 | 0.2923943 | 0.6444882 | 0.5092138 | 0.16039813 |
| Thorax width | 0.4885045 | -0.4260704 | -0.1915288 | 0.2381063 | -0.7027213 |
| Petiole width | 0.4839106 | -0.2114626 | 0.3745611 | -0.8026556 | 0.02599863 |
| Head length | 0.4235369 | 0.7381873 | -0.4726698 | -0.1544572 | -0.1023707 |
| Thorax length | 0.3637573 | -0.378593 | -0.4292307 | 0.1260703 | 0.68505439 |

**Table S6:** Tukey-corrected pairwise p-values comparing body size of adults emerging from treatments of early-stage embryos

Morphometric measurements were subjected to a principal component analysis followed by linear regression of principal component 1 scores. (EM=ergatoid male, QU=queen, WO=worker, Ethanol=solvent control, Methoprene=Methoprene)

| contrast | estimate | SE | df | t.ratio | p.value |
| --- | --- | --- | --- | --- | --- |
| QU Ethanol - WO Ethanol | 5.34767924 | 1.16591329 | 68 | 4.58668694 | <0.001 |
| QU Ethanol - EM Ethanol | 6.8213453 | 1.19811419 | 68 | 5.69340166 | <0.001 |
| QU Ethanol - QU Methoprene | 2.5353564 | 1.27719401 | 68 | 1.98509887 | 0.36149128 |
| QU Ethanol - WO Methoprene | 5.62962874 | 1.16591329 | 68 | 4.8285141 | <0.001 |
| QU Ethanol - EM Methoprene | 7.28148463 | 1.19315303 | 68 | 6.10272484 | <0.001 |
| WO Ethanol - EM Ethanol | 1.47366606 | 0.42996758 | 68 | 3.42738882 | 0.01276239 |
| WO Ethanol - QU Methoprene | -2.8123228 | 0.61694332 | 68 | -4.5584785 | <0.001 |
| WO Ethanol - WO Methoprene | 0.2819495 | 0.32977008 | 68 | 0.85498813 | 0.95569552 |
| WO Ethanol - EM Methoprene | 1.93380539 | 0.4159431 | 68 | 4.64920654 | <0.001 |
| EM Ethanol - QU Methoprene | -4.2859889 | 0.67582755 | 68 | -6.3418381 | <0.001 |
| EM Ethanol - WO Methoprene | -1.1917166 | 0.42996758 | 68 | -2.7716429 | 0.07458445 |
| EM Ethanol - EM Methoprene | 0.46013933 | 0.49913172 | 68 | 0.92187955 | 0.93954718 |
| QU Methoprene - WO Methoprene | 3.09427234 | 0.61694332 | 68 | 5.01548883 | <0.001 |
| QU Methoprene - EM Methoprene | 4.74612823 | 0.66699282 | 68 | 7.11571112 | <0.001 |
| WO Methoprene - EM Methoprene | 1.65185589 | 0.4159431 | 68 | 3.97135061 | 0.0023295 |

**Table S7:** Loadings of size measurements of adults emerging from treatments of early-stage embryos on principal components 1-5

| Trait | PC1 | PC2 | PC3 | PC4 | PC5 |
| --- | --- | --- | --- | --- | --- |
| Head length | 0.466633 | 0.16134472 | 0.4159195 | 0.74501599 | 0.02497802 |
| Head width | 0.4580136 | 0.15049684 | 0.5414776 | -0.6508546 | -0.1538678 |
| Petiolus width | 0.1470961 | -0.974121 | 0.1604896 | 0.01621195 | 0.02970913 |
| Thorax length | 0.5175076 | -0.0343465 | -0.5798608 | 0.03314526 | -0.6215809 |
| Thorax width | 0.5320017 | 0.035031 | -0.4145089 | -0.1413438 | 0.7671084 |

**Table S8**: Tukey-corrected pairwise p-values comparing log-transformed *krüppel-homolog* *1* expression measured in queen- and worker-destined late-stage embryos and larvae

(Q=queen, W=worker, Ctrl=handling control, EtOH=solvent control, JH=juvenile hormone III, Met=Methoprene)

| contrast | estimate | SE | df | t.ratio | p.value |
| --- | --- | --- | --- | --- | --- |
| Q Ctrl - W Ctrl | -0.2819789 | 0.39828281 | 42 | -0.7079867 | 0.99630065 |
| Q Ctrl - Q EtOH | -0.0049995 | 0.56776721 | 42 | -0.0088055 | 1 |
| Q Ctrl - W EtOH | -0.2869784 | 0.67811374 | 42 | -0.423201 | 0.99986869 |
| Q Ctrl - Q JH | -1.8928876 | 0.57182715 | 42 | -3.3102443 | 0.03690689 |
| Q Ctrl - W JH | -2.1748665 | 0.73741408 | 42 | -2.9493152 | 0.08798166 |
| Q Ctrl - Q Met | -1.2414213 | 0.5637081 | 42 | -2.2022414 | 0.37111319 |
| Q Ctrl - W Met | -1.5234002 | 0.73805731 | 42 | -2.0640677 | 0.4536454 |
| W Ctrl - Q EtOH | 0.27697948 | 0.70861793 | 42 | 0.39087281 | 0.99992304 |
| W Ctrl - W EtOH | -0.0049995 | 0.56776721 | 42 | -0.0088055 | 1 |
| W Ctrl - Q JH | -1.6109086 | 0.65379771 | 42 | -2.4639252 | 0.23891732 |
| W Ctrl - W JH | -1.8928876 | 0.57182715 | 42 | -3.3102443 | 0.03690689 |
| W Ctrl - Q Met | -0.9594423 | 0.63879843 | 42 | -1.5019485 | 0.80205083 |
| W Ctrl - W Met | -1.2414213 | 0.5637081 | 42 | -2.2022414 | 0.37111319 |
| Q EtOH - W EtOH | -0.2819789 | 0.39828281 | 42 | -0.7079867 | 0.99630065 |
| Q EtOH - Q JH | -1.8878881 | 0.54984324 | 42 | -3.4335025 | 0.02688223 |
| Q EtOH - W JH | -2.1698671 | 0.73503155 | 42 | -2.9520734 | 0.0874318 |
| Q EtOH - Q Met | -1.2364218 | 0.54202031 | 42 | -2.2811356 | 0.32759422 |
| Q EtOH - W Met | -1.5184008 | 0.73613745 | 42 | -2.0626593 | 0.45452094 |
| W EtOH - Q JH | -1.6059092 | 0.61777195 | 42 | -2.5995178 | 0.18480505 |
| W EtOH - W JH | -1.8878881 | 0.54984324 | 42 | -3.4335025 | 0.02688223 |
| W EtOH - Q Met | -0.9544429 | 0.60243844 | 42 | -1.5842994 | 0.7566709 |
| W EtOH - W Met | -1.2364218 | 0.54202031 | 42 | -2.2811356 | 0.32759422 |
| Q JH - W JH | -0.2819789 | 0.39828281 | 42 | -0.7079867 | 0.99630065 |
| Q JH - Q Met | 0.65146631 | 0.53040509 | 42 | 1.22824295 | 0.91873031 |
| Q JH - W Met | 0.36948736 | 0.67091526 | 42 | 0.55072135 | 0.99925122 |
| W JH - Q Met | 0.93344525 | 0.65558387 | 42 | 1.42383804 | 0.84109185 |
| W JH - W Met | 0.65146631 | 0.53040509 | 42 | 1.22824295 | 0.91873031 |
| Q Met - W Met | -0.2819789 | 0.39828281 | 42 | -0.7079867 | 0.99630065 |
